# Fine-scale landscape genomics show asymmetric patterns of gene flow for the invasive mosquito *Aedes albopictus*

**DOI:** 10.64898/2026.05.29.728763

**Authors:** Jessica Y. Ding, Emily MX Reed, Michael H. Reiskind, Martha O. Burford Reiskind

## Abstract

Mosquito-borne viruses like dengue, Zika, and chikungunya pose increasing health risks in the United States due to the expanding range of *Aedes albopictus*, a highly invasive mosquito species that now has a global distribution. *Aedes albopictus* thrive in artificial containers associated with anthropogenic land use, allowing populations to reach high numbers in urban and suburban environments. While the global spread of *Ae. albopictus* has been well-characterized, the effects of heterogeneous urban landscapes on dispersal and gene flow at fine spatial scales remain unclear. This study analyzed the genetic connectivity of *Aedes albopictus* populations collected in Wake County, North Carolina in 2018. We used single nucleotide polymorphisms SNP data from double-digest restriction-enzyme associated DNA sequencing (ddRADseq), and examined genetic connectivity through principal component analysis (PCA) and genetic network analysis. We then evaluated migration and source sink dynamics using a Bayesian approach for SNP data (BA3-SNP). We found little evidence of genetic clustering or isolated populations of *Ae. albopictus* in Wake County, suggesting high gene flow between sites. Migration analysis demonstrated asymmetric gene flow from rural to urban regions within Wake County, with greater gene flow occurring between and within urban regions. These findings suggest that *Ae. albopictus* populations demonstrate substantial gene flow within local metropolitan areas, with urban city centers serving as genetic sinks and surrounding suburban and rural regions serving as sources. This study highlights how heterogeneous landscapes shape mosquito population connectivity and migration at fine spatial scales, which is critical for informing vector control and public health intervention strategies.

## 1. INTRODUCTION

Understanding how organisms disperse across landscapes is a critical element of species ecology but remains a significant challenge. One approach applies population genetics in a spatially explicit context, which permits researchers to study the effects of heterogeneous landscapes on functional dispersal and gene flow (Aylward et al., 2020; Balkenhol, 2016; Fenderson et al., 2020; Klinga et al., 2019). Studies that connect population genetic approaches with movement across landscapes typically operate at fine spatial scales (tens to thousands of meters, associated with the movement ranges of the species) and are often used to inform local species and resource management. For example, previous research has used these approaches to investigate the effects of habitat fragmentation on metapopulation dynamics (Cheeseman et al., 2019; Yu et al., 2017), to evaluate the efficacy of landscape corridors and habitat restoration (Marquardt & Marcus, 2018; Sawaya et al., 2014; Soanes et al., 2018), and to manage the spread of zoonotic diseases (Biek & Real, 2010; Hemming-Schroeder et al., 2018; Richardson et al., 2017). Understanding the landscape features across natural and built environments that facilitate or inhibit gene flow is critical for managing invasive species, whose trajectories are often linked to human-induced climate and land use change.

*Aedes albopictus* is an invasive, cosmopolitan mosquito adapted to a range of climatic and anthropogenic environments (Hawley, 1988; Manica et al., 2016). Native to northern Asia, *Ae. albopictus* is now globally distributed and currently spreading through much of Europe and North America (Bonizzoni et al., 2013). Their rapid spread has raised international concern and increased monitoring and control of the species, as they are both a nuisance pest and a vector for viral diseases including dengue, Zika, and chikungunya viruses (Gratz, 2004; New York State Department of Health, 2025). Much of its introduction success can be attributed to its oviposition ecology and adaptation to anthropogenic landscapes. As a container-breeding species, *Ae. albopictus* females lay their eggs in small, ephemeral bodies of water. The eggs go through a drying and re-wetting cycle before hatching and can survive in both natural and artificial containers (Hawley et al., 1987). This ability has allowed *Ae. albopictus* to exploit international trade routes—for example, the shipping of used automobile tires—to move around the globe (Hawley et al., 1987).

Human-aided dispersal has been established as the primary driver of *Ae. Albopictus* movement across long distances, and has been shown to play a significant role in its introduction to non-native ranges (Gloria-Soria et al., 2022; Hawley et al., 1987; Medley et al., 2015; Schmidt et al., 2017). However, there is less certainty about the environmental and anthropogenic factors that mediate *Ae. albopictus* dispersal at finer spatial scales, particularly within urban landscapes. While mosquito control in the United States is often guided by local governments or individual homeowner decisions, many of the population genetics studies on *Ae. albopictus* and congeneric species have been conducted at regional, national, and continental scales. These studies have found that human transportation networks can drive long-distance dispersal and gene flow of container-breeding *Aedes* species beyond their natural dispersal capabilities (approx. 200m) (Gloria-Soria et al., 2022; Hopperstad et al., 2019; Medeiros et al., 2017; Medley et al., 2015; Schmidt et al., 2017). The movement of *Ae. albopictus* has been directly observed via passenger vehicles which likely exceed their natural dispersal distance (Eritja et al., 2017). Hopperstad et al. (2019) detected signals of directional migration out of the more densely populated urban center to less populated areas for *Ae. aegypti* populations along an interstate highway in Florida,, and Guagliardo et al. (2019) found boat traffic significantly explained gene flow of *Ae. aegypti* along the Amazon. These are some of only a few population genetics studies of *Aedes* mosquitoes that measured directional gene flow in heterogeneous landscapes, and we know of no fine scale landscape genetic studies of *Ae. albopictus*. One fine scale (1 to 10 km) population genetic study on *Ae. albopictus* in Wake County, NC found evidence of a temporally dynamic genetic structure that involved potential source and sink locations suggesting local dynamics as a driver (Reed et al., 2026). This was consistent with a study of *Ae. aegypti* within the city of Buenos Aires, Argentina, that uncovered more structure in less urbanized areas (Maffey et al., 2022). However, this previous study did not include the underlying landscape features, only broad-scale categories of impervious surfaces, leaving open the question of how landscape features affect gene flow. Furthermore, the previous study of *Ae. albopictus* in Wake County, NC did not directly measure fine-scale movement patterns that we address here.

Understanding how the underlying landscape drives population connectivity and directs migration will inform adaptive and effective mosquito management. For instance, releasing genetically modified mosquitoes to suppress vector populations and control arboviruses or understanding the distribution of insecticide resistant genes will benefit from understanding how genes move across the landscape (Abernathy et al., 2022; Dimopoulos, 2019; Rašić & Marshall, 2025; Richards et al., 2019; Unlu et al., 2016). To design an informative landscape genetics study, it is necessary to *a priori* define scales that are appropriate for the species under study, the genetic data available, the landscape variables of interest, the computational intensity for geospatial data, and the level at which management operates(Cushman & Landguth, 2010). If the ecological and landscape processes governing *Ae. albopictus* dispersal are scale-dependent, findings from larger regional landscape genetics studies may not be appropriate for local control. For instance, if the scale of a population genetic study is at the level of the southeastern United States, it may obfuscate the geographic pattern of insecticide resistant alleles within smaller, county-level spatial scales.

To provide insight into how this important vector species moves across urban landscapes, we investigated the spatial genomic patterns of *Ae. albopictus* within Wake County, North Carolina. This study uses previously sampled and analyzed population genomic data for Wake County (Reed et al., 2026) to describe site to site patterns of gene flow. Here, we use a network approach as well as a Bayesian landscape genomic analysis that allows us to test the hypothesis that *Ae. albopictus* mosquitoes use human movement pathways to disperse across the landscape from sources to sinks, consistent with larger scale studies of *Aedes spp.* population genetics (Gloria-Soria et al., 2022; Hopperstad et al., 2019; Medley et al., 2015; Schmidt et al., 2017). In this study we refer to a source as a location with greater emigration than immigration and a sink as a population with greater immigration than emigration. From this and knowledge that *Ae. albopictus* is an anthropophilic vector typically found in intermediate or low human densities (Reiskind & Lounibos, 2013), we predict within Wake County we will observe (1) higher genetic connectivity among areas with higher human density, and lower genetic connectivity among areas with lower human density, and (2) greater gene flow from lower human density (source) areas to higher human density areas (sink).

## 2. MATERIALS AND METHODS

### 2.1. Study system

Our study was within a 30 minute drive of our facilities for logistical reasons, with all sites within the political boundary of Wake County, NC. Mosquitoes, including *Aedes albopictus*, have been collected annually in at least some locations in the county since 2015 (Reed et al., 2026; Spence Beaulieu et al., 2019). Wake County is a rapidly urbanizing area—between 2010 and 2018, its population grew by 20.3% from 906,882 to 1.091 million residents, making it representative of the rapid growth seen in the region (Terando et al., 2014; U.S. Census Bureau, n.d.). Wake County is home to Raleigh, the state’s capital and second largest city with an estimated population of 457,159 in 2018, when the samples for this study were collected (U.S. Census Bureau, n.d.). Surrounding the major urban center of Raleigh is a mosaic of land use types, including suburbs, agriculture, and preserved open and forested green spaces (City Planning, 2025; Terando et al., 2014). This rural/suburban transition has spread outward from the Raleigh epicenter to towards the outskirts of the county in a pattern consistent with descriptions of urbanization in the southeast (Terando et al., 2014). The areas west of Raleigh—including the Apex, Cary, and Morrisville municipalities—are the most developed region of Wake County, with the eastern and southern portions of the county slated for significant residential and mixed-use development in the next two decades (City Planning, 2025; *PLANWake Comprehensive Plan*, 2021). All regions were assigned a classification of a broad Urban or Rural designation (Suppl. **Table S1, Figure S1**).

Wake County does not conduct mosquito surveillance or control regularly, and mosquito adulticiding is entirely privatized(Wake County Public Health,, retrieved March 28, 2026)). We used the previously published data of 61 sites where *Ae. albopictus* were collected within Wake County from June 7 to June 25, 2018, using Biogents BG sentinels baited with BG Lures ™, a chemical attractant designed for *Ae. albopictus* and *Ae. aegypti* (Biogents GmbH, Regensburg, Germany; Reed et al., 2026; **Figure 1A**). To identify sites, Reed et al. (2026) used a random point generator using the *r.random.cells* function in GRASS GIS (Neteler et al., 2012) to select 100 points across Wake County with a minimum distance of 1000m between points (<1km dispersal distance; Honório et al., 2003). For each point, Reed et al. (2026) determined whether the site was accessible by examining the landscape within a 100m buffer of the point using Google Earth (Google, 2008). A site was deemed inaccessible if its buffer was entirely water or on private land. In addition, sites needed to be within 1 km of a public road to minimize the amount of on-foot travel required to reach the sampling location. These requirements eliminated 27 locations, and they randomly selected 60 of the 73 remaining sites using the *sample.int* command in RStudio (RStudio Team, 2018; Reed et al., 2026). Four of the 60 sites were located on private roads not marked on Google Earth and were removed, leaving 56 randomly sampled sites in addition to five sites used for on-going surveillance for a total of 61 sites (described in Reed et al., 2019).

**Figure 1.**
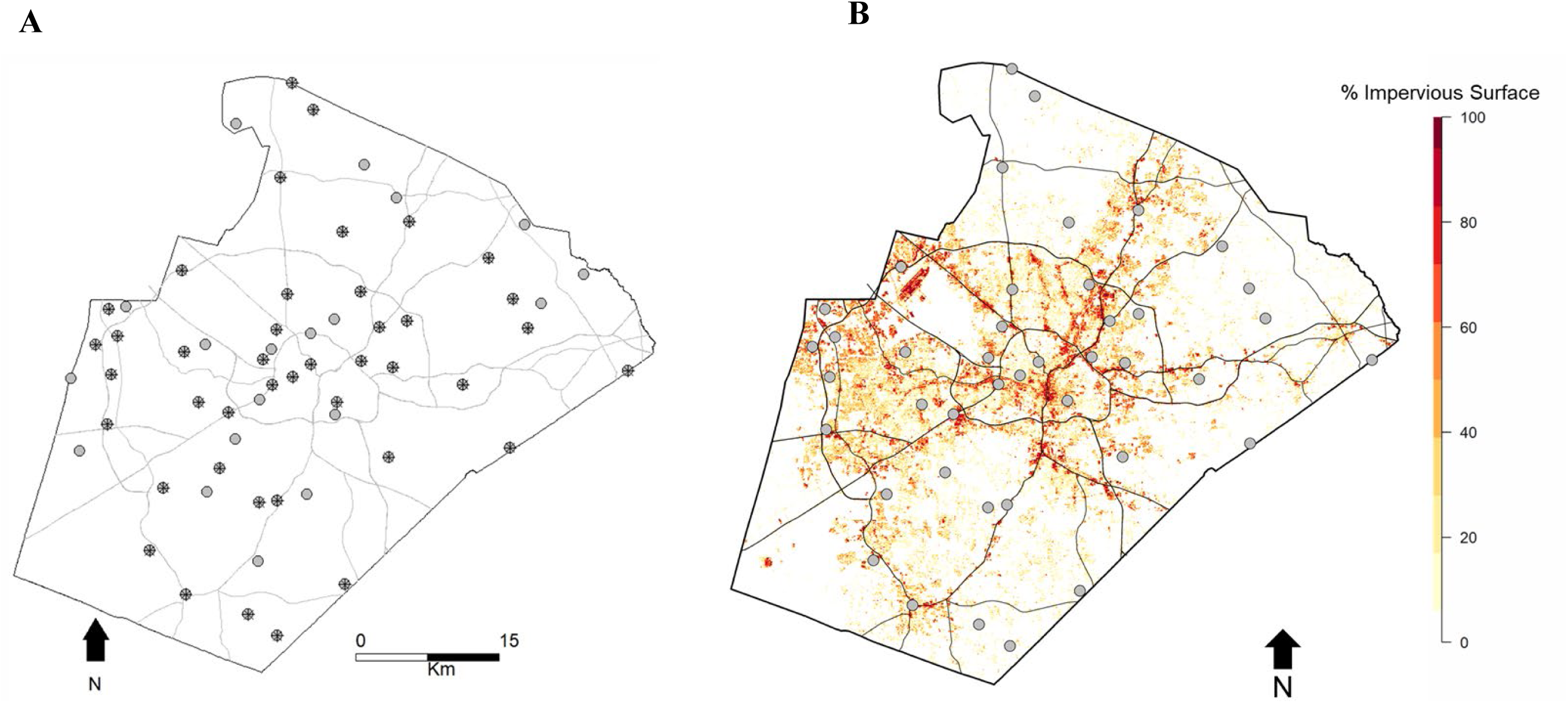
**A.** Wake County sampling locations (n = 61). Points overlaid with asterisks are sites that were retained for genomic analyses (n = 42). **B.** Sample sites for genomic sequencing in Wake County, North Carolina. Background shows percent impervious surface from the USGS 2016 impervious surface dataset and highways in Wake County. Raleigh is in the center of the county.

Locations were sampled once a week for three weeks, leaving traps out to collect mosquitoes for approximately 24 hours. After collection, Reed et al. (2026) identified all individuals collected and separated and sexed *Ae. albopictus* placing each individual into a microcentrifuge tube of 95% ethanol to preserve before DNA extraction (Suppl. **Table S2**).

### 2.2. DNA extraction and genomic library preparation

*Ae. albopictus* were identified at 59/61 sampled sites, with six or more individuals collected at 47 sites. DNA was extracted from *Ae. albopictus* individuals using the Qiagen DNeasy Blood & Tissue Kit (Qiagen Inc., Valencia, CA, USA) and DNA concentrations were quantified using a Qubit 2.0 fluorometer (Invitrogen, Carlsbad, CA, USA). After extraction of individual mosquitoes, there were high-concentration DNA samples (>8 ng/ul) for at least six individuals at 42 sites (**Figure 1B**), and sequence data per individual was obtained from 289 adult *Ae. albopictus*. Genomic DNA of individual mosquitoes were sequenced from sampled sites using double-digest restriction-enzyme associated DNA sequencing (*ddRADseq*) following Burford Reiskind et al. (2016) and as outlined in Reed et al. (2025). Each individual was given its own unique barcode and then sequenced (Suppl **Table S2**; Burford Reiskind et al., 2016; Reed et al., 2026).

### 2.3. Quality control and filtering

We used sequence data from the previously published study (Reed et al., 2026), which included sequence quality control and trimming and a catalog of SNPs using the STACKS *de novo* pipeline 1.09 (v.1.09; Catchen et al., 2011). For our landscape genetic study, we conducted a more conservative filtering of the data after the *de novo* pipeline than the previous study (see Reed et al., 2026). Here we filtered SNPs from the STACKS catalog using a *populations* pipeline that included SNPs present in at least 75% of individuals in a sample location and occurred in at least two sites. We further filtered SNPs by removing individuals with over 25% missing data and variants with a minor allele frequency (MAF) less than 0.01 and a genotyping rate of less than 0.75 in PLINK v1.19 (Purcell et al., 2007). We continued with a more stringent genotyping rate to accommodate for the sensitivity to missing data in landscape genetic and connectivity analyses, an issue that has been previously discussed regarding ddRADseq data (Arnold et al., 2013; Gautier et al., 2013; Schmidt et al., 2021; Sherpa et al., 2018). We converted PLINK files to Genetix files using PGDSpider v2.1.1.0 (Lischer & Excoffier, 2012) and imported these files to RStudio. With the *hw.test* function in the R package *pegas* v.0.14 (Paradis, 2010), we removed SNPs that were not in Hardy-Weinberg Equilibrium after the conservative Bonferroni correction (*P* < 0.05/# SNPs). We conducted a second suite of filtering steps in the R package *adegent* v 2.1.11 (Jombart, 2008) including removing individuals having greater than 25% missing data, removing populations with fewer than three individuals, and removing all loci with only one allele (**Suppl Table S3**). Finally, we converted the filtered file to a *locus* object as implemented in the R package *gstudio* (Dyer, 2016). We used this method because it allowed us to identify locus objects as data frames that we could easily summarize, manipulate, and transform the marker-based genetic data and integrate genetic and spatial data for downstream analyses.

### 2.4. Genetic structure and connectivity

We used principal component analysis (PCA) and pairwise *F_ST_* to assess genetic structure and differentiation between sites with this **new filtered data set**. In the previous study, pairwise comparisons on the larger SNP data set were conducted between sites; here we conducted PCAs at both the individual level and at the population level using the *dudi.pca* function in the R package *ade4* (Dray & Dufour, 2007). Prior to conducting PCA, we scaled and centered allele frequencies, with missing data replaced by mean allele frequencies at a given locus (Zaborowska et al., 2021). Only the first three axes were retained. To assess genetic differentiation between populations, we estimated pairwise *F_ST_* using the *hierfstat* package in R. We tested significant differences in pairwise *F_ST_* using 1000 bootstraps and generated 95% confidence intervals (CIs; boot.ppfst). We determined significant differences when CIs did not include zero up to four significant figures.

We evaluated genetic connectivity with a graph theoretic approach following Dyer and Nason (2004). Graph theory has been applied to study complex social and biological systems, where vertices, or nodes, represent elements of a given system, and edges represent the interactions or relationships between them (Barabási & Albert, 1999; Girvan & Newman, 2002). In the context of landscape genetics, individuals or sample sites can be conceptualized as a set of nodes connected by gene flow (Garroway et al., 2008). In contrast to traditional approaches such as *F-*statistics, a population graph uses a multivariate network model to define connections among groups of populations simultaneously (Savary et al., 2021b). This allows us to address multiple population genetic questions at once, such as genetic diversity, differentiation, and connectivity, to provide novel insights into how genes move across networks.

Population graphs are composed of a set number of nodes (in this case, the sites sampled in Wake County) and edges that connect nodes. We built population graphs using the R package *graph4lg* (Savary et al., 2021b), which expands upon the functionalities of Dyer and Nason’s *popgraph* package (2004). Starting with a fully saturated graph where all nodes are connected by edges, we pruned the edges using the conditional independence principle (CIP), which retains links between populations connected to each other based on the covariance of their allele frequencies (Savary et al., 2021b). This method removes the maximum number of edges while still accurately describing the among-population genetic variation (Dyer & Nason, 2004). The CIP method results in a pruned graph where the edges represent links between populations that are directly exchanging migrants through single generation dispersal events (Dyer & Nason, 2004; Savary et al., 2021b). Edges are weighted such that a larger edge weight represents greater genetic distance between two linked nodes.

Using the defined population graph, we calculated node-based metrics to evaluate genetic connectivity between sites. Node centrality measures can be used to identify influential nodes in a network. In the case of genetic networks, these are populations that play an important role in maintaining gene flow (Rodger et al., 2018). We calculated the following metrics for each site: degree centrality, betweenness centrality, closeness centrality, and mean inverse edge weight (M_IW_). Degree centrality is a count of the number of edges connected to a given node, where sites with low degree values are more genetically isolated (Newman, 2016). Betweenness centrality measures the extent to which a node lies on the shortest path between two nodes in a network, given all possible paths within the network (Koen et al., 2016; Newman, 2016). A node with higher betweenness centrality plays a more important role in regulating gene flow through the network, often acting as either a bridge or a bottleneck (Garroway et al., 2008). Closeness centrality is the reciprocal of the sum of the shortest path between a given node and all other nodes (Fournier et al., 2024; Koen et al., 2016). A node with low closeness centrality is, on average, a shorter distance from other nodes within the network—in other words, nodes with low closeness centrality are more genetically similar to other nodes in the network, while nodes with higher closeness centrality are more genetically distant from other nodes in the network (Newman, 2016). Mean inverse edge weight is the mean of the inverse weights of edges connected to a node. Because a larger edge weight represents greater genetic distance, nodes with higher *M_IW_* are more strongly connected to other neighboring nodes (Fournier et al., 2024). When considered together, these node-based network metrics can help identify hubs of gene flow and putative locations of genetic sources and sinks.

### 2.5. Gene flow and source sink dynamics

To assess source and sink dynamics, we measured gene flow between sites using the program *BA3-SNPs*, which uses the patterns of SNP data in a Bayesian framework to assess demographic independence within and among populations (Mussmann et al., 2019). We conducted five runs in *BA3-SNPs* with 5,000,000 iterations each, a burn-in period of 1,000,000, and an interval of 1000 iterations between samples. We tested for model convergence by comparing the parameterized migration rates between runs. Only migration rates with a 95% confidence interval greater than zero were included in the results. We evaluated source/sink dynamics by subtracting the sum of immigration rates from emigration rates following Gustafson et al. (2019). Higher emigration indicates the region is a genetic source, while higher immigration indicates a genetic sink.

Due to the computationally intensive process of estimating unidirectional migration rates and the rapid increase in pairwise calculations necessary with additional sites, we grouped sites geographically into ten smaller regions, each with three to five sites (**Figure 2A**). We assigned sites to regions based on their proximity to Wake County municipalities and major roadways. These regions are at different stages of urbanization (defined by % impervious surface, see **Supplement 1**), with the most urbanized areas in Raleigh and Cary and the least developed areas in the northwest and eastern parts of the county (**Figure 2B**).

**Figure 2.**
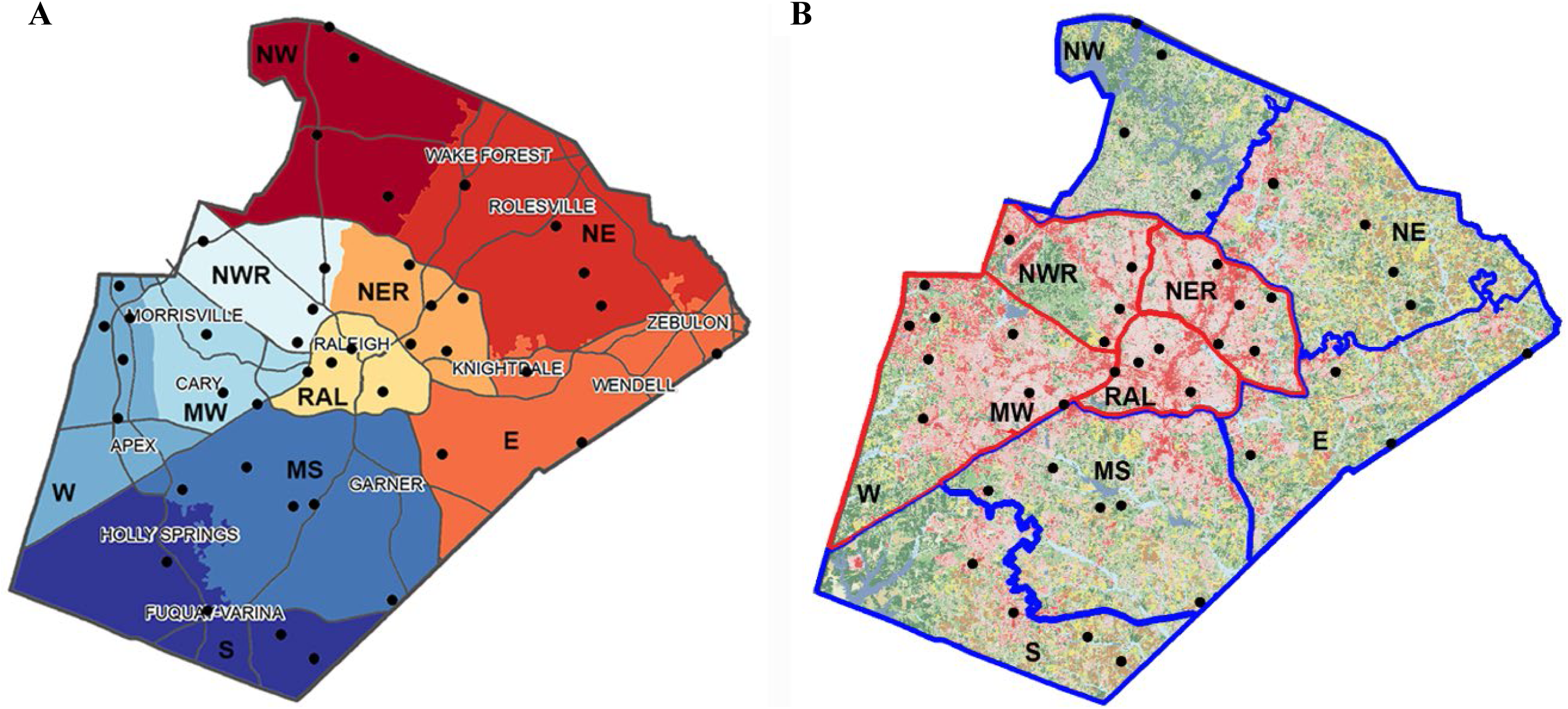
**A.** To compare node parameters and quantify directional gene flow, we grouped sample sites in ten regions of 3-5 sites. We determined sites based on proximity to Wake County jurisdictions (black labels with white background) and major road networks. Abbreviations for regions are shown in black: RAL = central Raleigh; NWR = northwest Raleigh; NER = northeast Raleigh, E = east; MS = mid-south; S = south; W = west; MW = mid-west; NW = northwest, and NE = northeast. **B.** Developed regions of Wake County outlined in red (W, MW, NWR, NER, RAL); less developed or rural regions outlined in blue (S, MS, E, NE, NW). Urban and rural categories were defined based on percent impervious surface (red pixels) (see **Supplement 1**).

## 3. RESULTS

### 3.1. Quality control and filtering

As the previous STACKS *de novo* pipeline identified 3,096,027 SNPs (Reed et al., 2026), the post-*population* pipeline for this study retained 183,753 SNPs. PLINK filtering removed 179,704 SNPs and 44 individuals that did not meet thresholds for minor allele frequency and genotyping rates (Suppl **Table S3**). For this study, we minimized missingness and loci that are out of HW equilibrium or linked for a final, conservative dataset of 245 individuals and 4,013 loci across 39 sites.

### 3.2. Genetic structure and connectivity

At the population level, PC1 and PC2 accounted for 21.59% and 17.03% of the variance in allele frequencies, respectively (**Figure 3**). Collectively, the first three principal components accounted for 53% of the observed variance in the SNP data. The majority of sites across Wake County formed a single cluster, with populations from all ten regions represented within the cluster (**Figure 3, Suppl. Figure S2**). While a few sites were visibly differentiated from the main cluster in the PCA (S04, S11, S27, S48), these sites were also distant from each other and did not demonstrate clear clustering patterns (**Figure 3, Suppl. Figure S2)**. Within the main cluster, many of the sites in close proximity were from different regions of Wake County, which suggests that major roadways may not be a barrier to gene flow. Based on the PCA results, mosquito populations in Wake County demonstrated genetic clustering, with a few sites being more genetically differentiated compared to the others.

**Figure 3.**
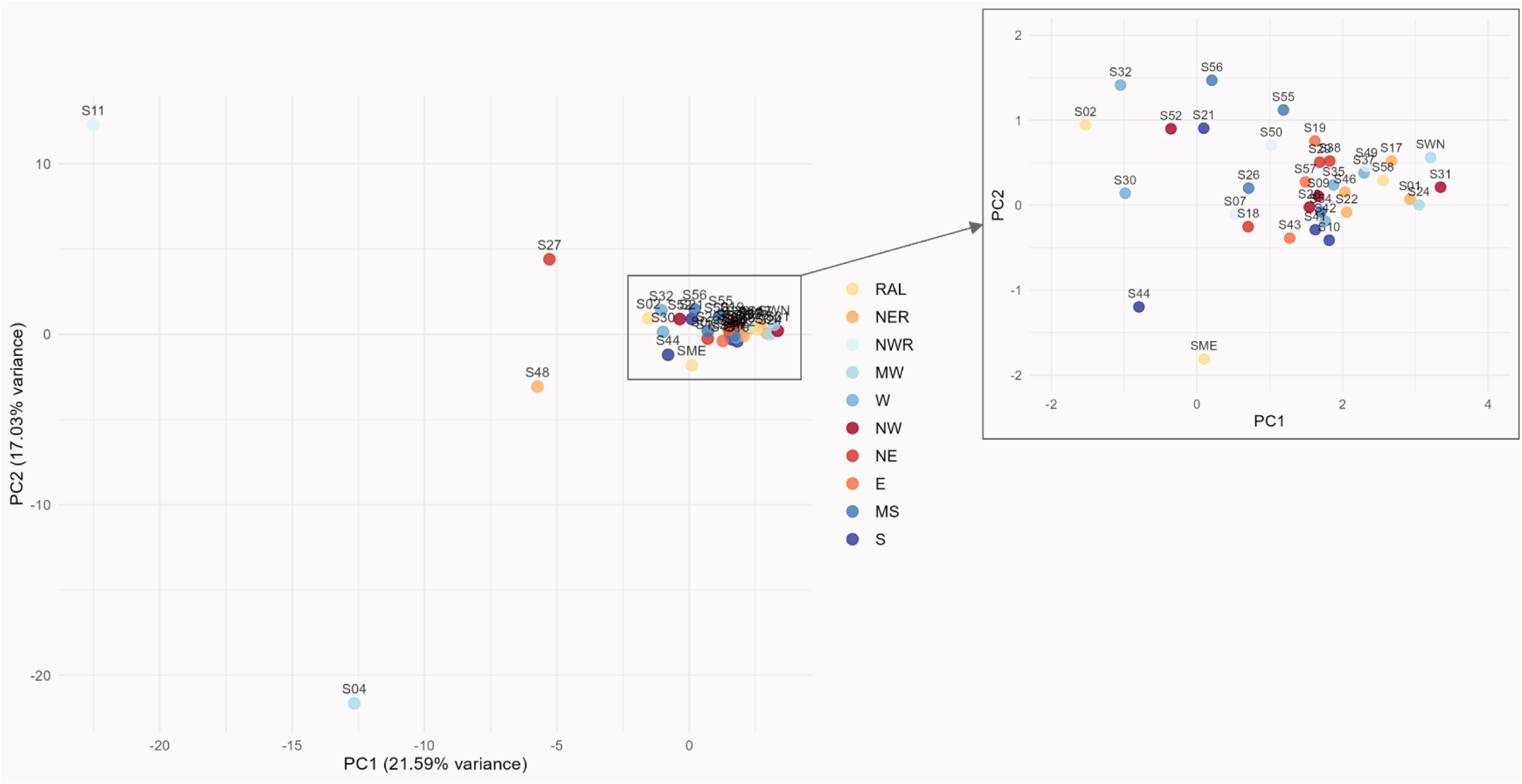
Population-level PCA with PC1 and PC2 on the *x* and *y* axes, respectively. Points are colored according to Wake County regions (see Figure 2). The first two principal components account for 38.62% of the variance in the dataset. The majority of points form a single cluster, representing sites that have greater genetic similarity. S04, S11, S27, and S48—all located in urban regions—are separated from the main cluster, indicating that these sites are more genetically differentiated from the rest of the sites in Wake County. The zoomed in portion of the plot reveals some clustering by region (e.g. E, NER), but no clear patterns.

The pairwise *Fst* analysis showed similar patterns of genetic structure ranging from – 0.0001 to 0.0179 (**Table 1**). The sites with the greatest number of significant pairwise differences were found in the more developed Urban regions (RAL, NER, NWR, MW, and W; Fig. 2B), which had 21.86% significant pairwise comparisons out of the total pairwise comparisons. In contrast, the less developed Rural regions (NW, NE, E, MS, S; Fig 2B) had significant pairwise comparisons in only 12.96% of the comparisons, a significant difference between Urban and Rural broad regions (binomial test, *Z* = 4.55, p < 0.001). For sites within the broad Urban region, 56.79% of the significant differences were with other urban sites. Individual sites with the greatest numbers of significant pairwise *Fst* values were S24, S11, S26, S50, and S37, with S24 having the highest at 21 significant differences. Four out of these five sites were in Urban regions: S24 in midwest Wake County (MW), S11 and S50 in northwest Raleigh (NWR), and S37 in western Wake County (W) (see **Figure 2B**). S26 was in mid-southern Wake County (MS), which was classified as Rural. With the exception of the MS region that had genetic differentiation within, mosquito populations in Rural regions did not demonstrate significant genetic differentiation from other sites within their respective smaller regions. Furthermore, three sites located in rural NE and E regions demonstrated no significant differences with any of the sites sampled in Wake County (S29, S38, and S57).

**Table 1.**
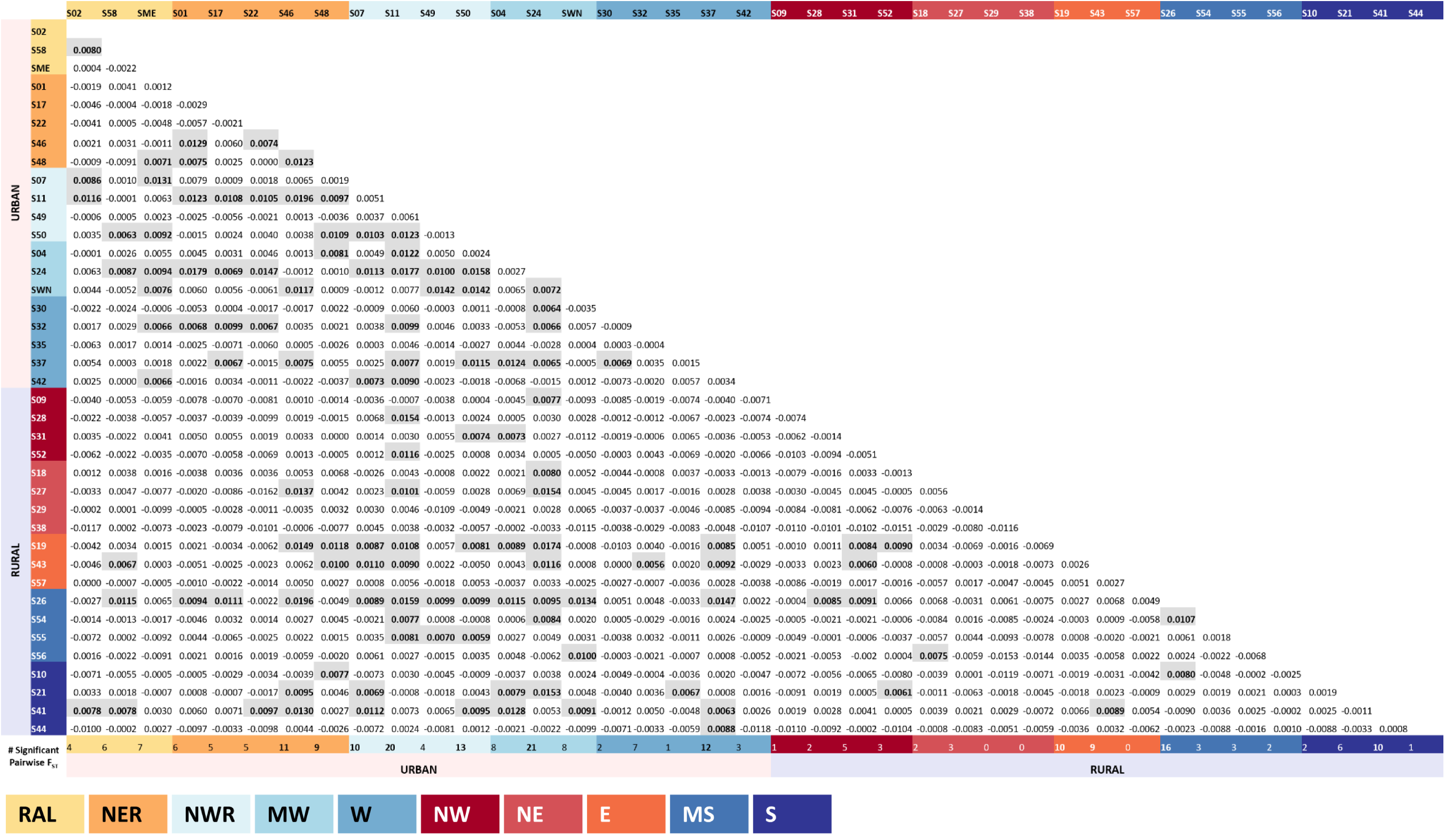
*Aedes albopictus* pairwise *F_ST_* values among sites in Wake County 2018 using the R package *hierfstat*. Bolded numbers with gray shading are comparisons that were identified as significantly differentiated following a bootstrap analysis in *hierfstat* (bootstraps = 1000) generating 95% CIs. We restricted our significance test using CIs that did not include zero to three significant figures. Background color of site names separated by region with the key in the caption and RAL, NER, NWR, MW, and W more urban regions and NW, ME, E, MS, S more rural regions. The last row in the number of significant pairwise comparisons for each site by the column header.

The final population graph consisted of 39 nodes connected by 35 edges, with 7 isolates—populations with no connections to other nodes based on the parameters in our analysis (**Figure 4A**). Degree centrality ranged from zero to five, with S58 and S24 having the highest number of connections (**Table 2**). These sites were in central Raleigh (RAL) and midwest Wake County (MW), respectively. On average, sites in MW, NER, and NW had the highest degree centrality, representing greater connectivity, while sites in NE, NWR, and S had the lowest degree centrality, indicating low connectivity with other sites. Betweenness centrality ranged from 0 to 180, with higher betweenness centrality indicating a more central role in maintaining gene flow through the network. S24 in MW had the highest betweenness centrality, followed by S58 in RAL, S46 in NER, and S31 in NW (180, 161, 101, and 98, respectively). On average, sites in MW and RAL had the highest betweenness centrality, which suggests that these sites serve as “hubs” that are critical in maintaining gene flow across the network. In contrast, sites in NWR and NE had the lowest average degree centrality and betweenness centrality, meaning sites in these regions are more genetically isolated relative to the rest of the county. Closeness centrality measures the mean distance from a given node to all other nodes in the network (Newman, 2016). Unlike the previous centrality measures, closeness centrality is interpreted such that lower values indicate greater genetic relatedness (Fournier et al., 2024). Mean closeness centrality was lowest in east and west Wake County and highest in south Wake County and northwest Raleigh—in other words, sites in E and W were more genetically related to mosquito populations across Wake County, while sites in S and NWR were the most genetically distant among all mosquito populations.

**Figure 4.**
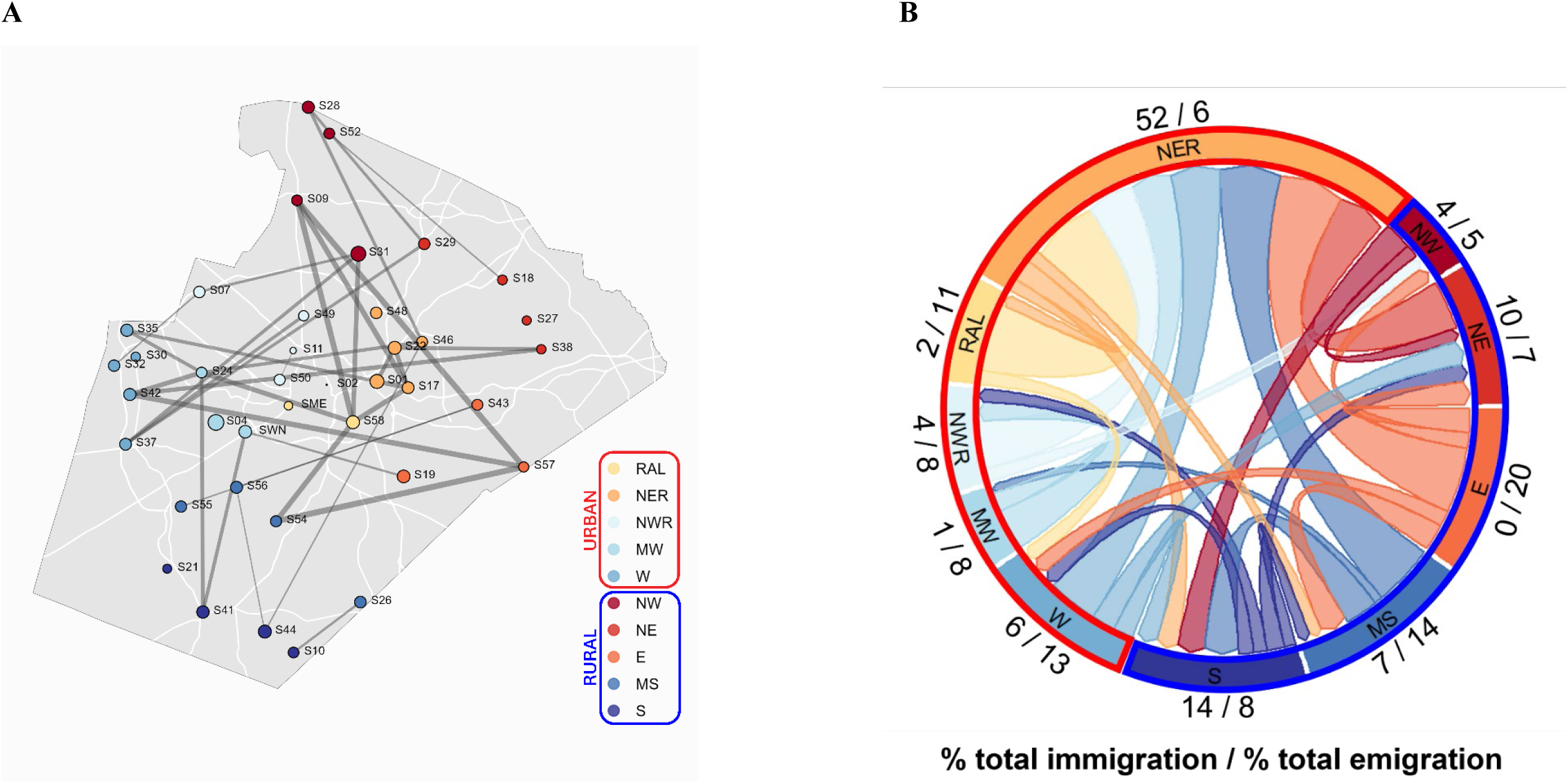
**A.** Population graph generated using the *graph4lg* package, with nodes positioned based on their geographic coordinates. Node size represents *H_O_*, the observed heterozygosity at each sampled site, and node color indicates Wake County region. Nodes are connected by edges that represent gene flow between sites—for ease of interpretation, edge thickness in this figure is proportional to the inverse genetic distance—therefore, thicker edges represent greater genetic connectivity and thinner edges represent less genetic connectivity. Isolates, or nodes without any edges, represent genetically isolated sites. **B.** Migration between Wake County regions estimated by *BA3-SNPs*. Width of arrows indicates migration rate, and fractions along outside of regions indicate the percent of relative immigration and the percent of relative emigration moving into and out of the region, respectively. Locations outlined in blue are rural and red are urban.

**Table 2.**
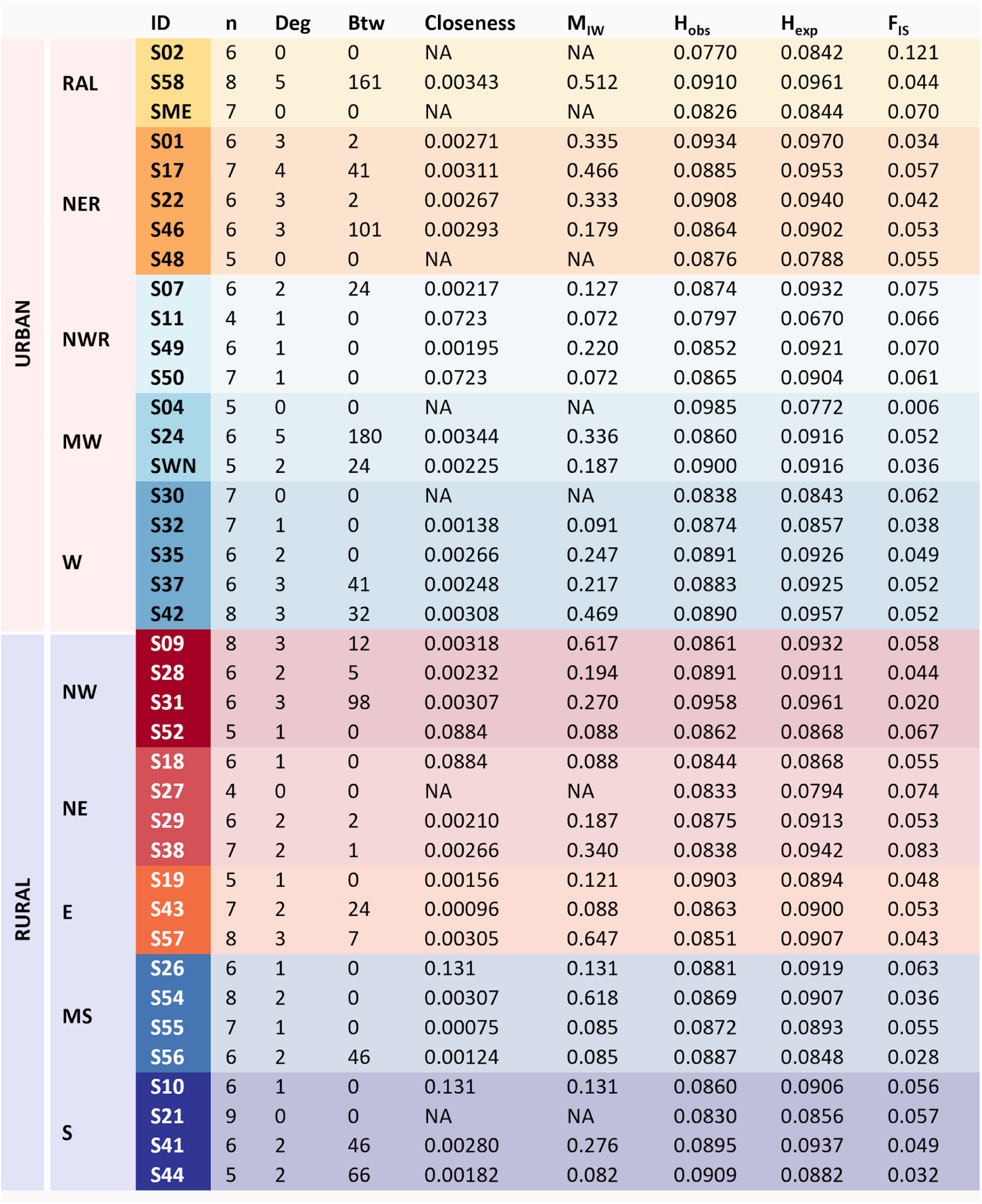
Genetic network summary statistics for the population graph created using the R package *graph4lg*. Of the 39 sites (nodes), 7 were isolates with no connections to other nodes, while the rest were connected by 35 edges. We calculated degree centrality (deg), betweenness centrality (btw), closeness centrality (closeness), and mean inverse edge weight (M_IW_). Observed heterozygosity (H_obs_), expected heterozygosity (H_exp_), and inbreeding coefficient (F_IS_) were calculated by Reed et al. (2025).

While the node centrality measures described above are calculated based on the number of connections a node has (i.e. the presence or absence of an edge), mean inverse edge weight takes into account the degree of genetic relatedness between nodes (Koen et al., 2016). Because a larger edge weight represents greater genetic distance, a site with higher mean *inverse* edge weight is *more closely related* to the nodes it shares a connection with. The five sites with the highest *M_IW_* were S57, S54, S09, S58, and S42. These sites were strongly connected to each other despite being in different regions of Wake County (Fig. 5A). The most central regions (RAL and NER) had the highest *M_IW_* (0.512 and 0.329, respectively), while NWR and S had the lowest *M_IW_* (0.123 and 0.163, respectively) which is consistent with low gene flow both into and out of these more peripheral regions. Overall, when we compared node characteristics between regions defined *a priori* (Figure 2A), we found the strongest patterns of genetic connectivity in central Raleigh (RAL) and the regions immediately surrounding the city center (NER, MW, NW).

**Figure 5.**
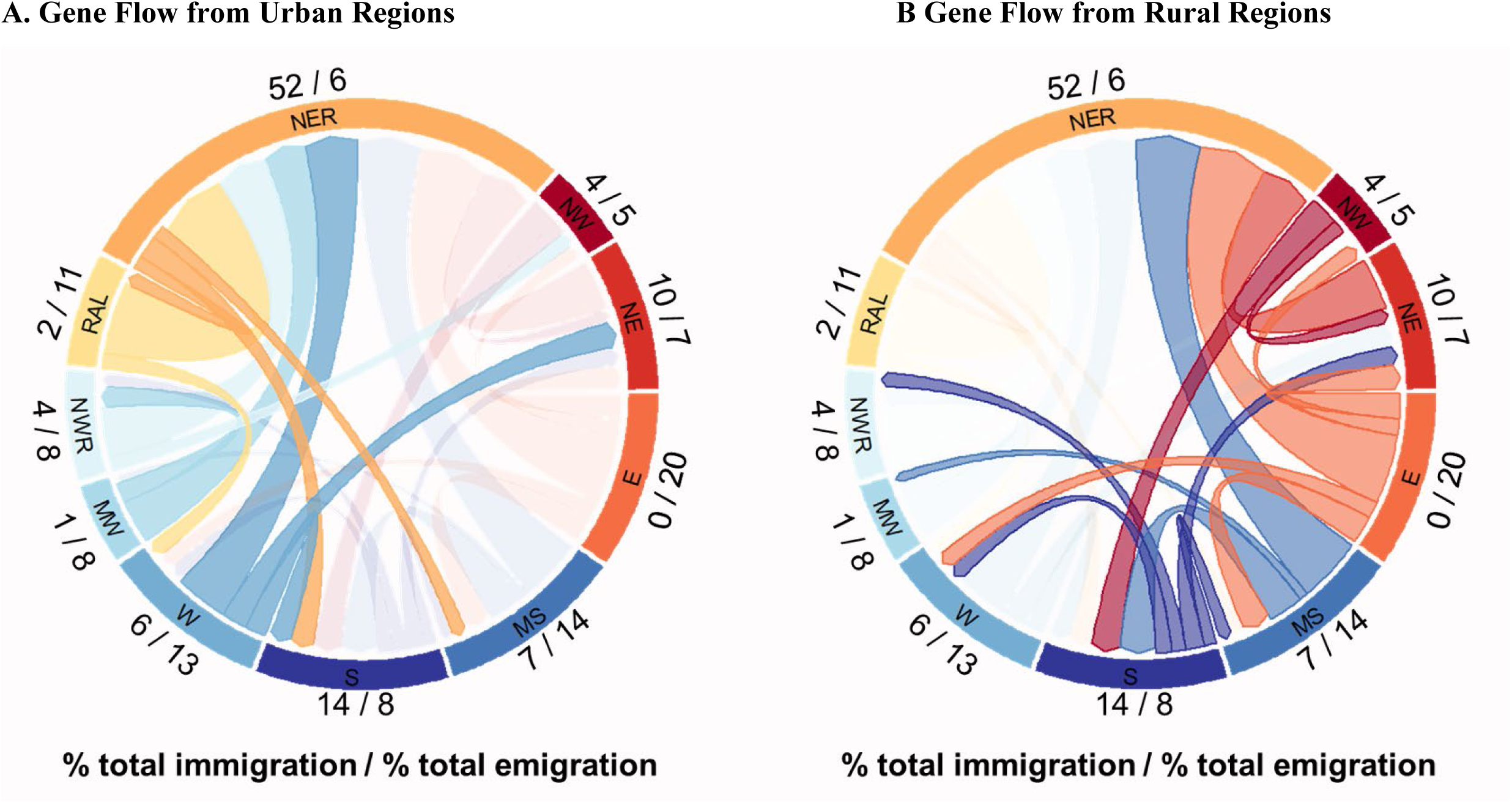
Directional gene flow between regions of Wake County. Only estimates of migration that had a 95% confidence interval that did not include zero are included. Width of arrows indicates migration rate, and fractions along outside of regions indicate the percent of relative immigration and the percent of relative emigration moving into and out of the region, respectively. **A. Gene flow from Urban regions** - highlights gene flow originating from urban regions of Wake County, while **B. Gene flow from Rural regions** - highlights gene flow originating from rural region.

We found similar genetic diversity (*H_E_*) and inbreeding coefficients (*F_IS_*) at individual sites with the exception of four sites with slightly lower genetic diversity in the Urban landscape (SO2 in RAL, S48 in NER, S11 in NWR, and SO4 in MW) and one site in the Rural landscape (S27 in NE) (**Table 2**). We also found one site with elevated *F_IS_* in the Urban landscape (S02 in RAL).

### 3.3. Gene flow and source sink dynamics

Of the 90 estimated migration rates between sites, we found 27 that had 95% confidence intervals above zero (**Figure 4B**). Northeast Raleigh (NER) had an immigration rate more than eight times greater than its emigration rate and was estimated to receive 52% of total emigrants from both urban and rural regions (**Figure 5**), making it a net genetic sink. The only regions that did not have statistically significant emigration rates into NER were the northwest (NW) and southern (S) regions. S, along with NE, were also identified as potential genetic sinks due to higher rates of immigration compared to emigration. In contrast, rates of gene flow into and out of northwest Wake (NW) were approximately equal. The largest source population was from the rural eastern region (E), which contributed 20% of all emigrants and did not receive any immigrants. Central Raleigh (RAL) and eastern Cary (MW) were also identified as potential source populations, with gene flow primarily moving from those regions into NER. While NWR, W, and MS received some immigrants, their emigration rates constituted a higher proportion of their total migration gene flow.

## 4. DISCUSSION

We found mosquito populations located in the more developed, urban regions of Wake County demonstrated greater genetic differentiation compared to sites in rural regions (PCA and pairwise *F_ST_*). Sites with the greatest genetic differentiation were in regions immediately adjacent to the urban centers of downtown Raleigh or downtown Cary and Morrisville (NER, MW, W), while sites with the least genetic differentiation were in rural regions on the periphery of Wake County (NW, E, S). The patterns we observed are consistent with multiple introductions of a few individuals of an invasive species into distinct sites in an urban environment. When multiple introductions occur, each individual site will have reduced genetic diversity and have unique allelic combinations, which can cause an increased degree of genetic divergence in pairwise site comparisons (Reed et al., 2020). While *F_ST_* can be helpful in evaluating genetic differentiation at this scale, it assumes migration-drift equilibrium, which is not always the case when gene flow is high (Savary et al., 2021a). The previous study examined patterns of *F_ST_* and *F_IS_* in this landscape, finding only a significant difference between urban and rural locations in inbreeding coefficients (*F_IS_*), with lower genetic diversity and higher *F_IS_* in the central, urban area of Raleigh (designated “RAL” here) (Reed et al., 2026). While this is consistent with what we expected, it was limited in scope because pairwise estimates treat all population pairs as equally weighted (Richardson et al., 2016). Our network approach in this study improves the sensitivity of our landscape genetic analyses because we can examine the patterns of gene flow between sites (nodes) while accounting for the genetic covariance among all populations, giving more insight into how mosquitoes move around the landscape (Dyer et al., 2010; Richardson et al., 2016).This allows us to address gene flow among sites within broad regions of urban and rural, sites that function as connectors or “hubs”, source and sink dynamics, and directionality of gene flow.

Our combined approach of using both a network and migration analysis provided a more comprehensive understanding of the complicated gene flow dynamics of *Ae. albopictus* in Wake County, NC, which includes identifying hubs or connecting sites, and sites that are more or less isolated. While our migration analysis aggregated sites by region, which could make finer-scale patterns harder to differentiate site-specific metrics from the network analysis allowed us to tease apart these details. This showed that genetic connectivity varied between regions of Wake County, with certain sites acting as genetic sources or sinks. These results combined with the results from the pairwise *F_ST_* analysis showed that one site S24 in MW was the most genetically differentiated and played a central role in the genetic network (highest betweenness centrality and high *M_IW_*). With high values of *M_IW_* that indicate high immigration, high emigration, or both (Koen et al., 2016), a site would be genetically differentiated due to high rates of admixture between unrelated individuals migrating into the site. Therefore, S24 most likely served as a central “hub” through which mosquito dispersal occurred. In general, sites with the highest betweenness centrality values were in the most developed urban centers in Wake County—central Raleigh and Midwest Wake, which together encompass the major urban municipalities. These sites were highly influential in maintaining connectivity between these regions, supporting our prediction of greater genetic connectivity among more urban, developed areas in Wake County.

The combined network and migration analyses suggested sources and sinks in Wake County. Specifically, with the highest *M_IW_* on average, RAL and NER, represent these putative genetic sources or sinks. Our migration results also showed NER as the single largest genetic sink, and RAL and MW as significant sources going into NER. While the Rural southern region (S) had low degree centrality, it had a relatively high mean betweenness centrality, which suggests that although these sites may not be well-connected, they nevertheless serve as important “bridges” connecting rural and urban regions. These results support our prediction of greater gene flow from more rural to highly developed urban regions of Wake County.

Our migration results showed a general trend that genetic material flows from rural to urban regions and between urban regions, consistent with human aided dispersal, a phenomenon noted for this species (Eritja et al. 2017). The largest genetic sink was in urban northeast Raleigh (NER), which contributed only six percent of immigrants but received 52% of the total migration across the county. The less developed eastern region was the greatest single source of gene flow, with almost half of the emigration from this region (E) flowing into urban Raleigh (NER). Our migration rates showed highly asymmetric gene flow. Overall, migration into urban regions constituted the majority of gene flow. In addition, migration originating from urban regions flowed primarily into other urban regions in the county, with urban-to-rural migration representing the smallest proportion of observed gene flow. Taken together, our network analysis demonstrated strong genetic connectivity across urban and rural regions of Wake County, while our migration analysis showed asymmetric gene flow from rural regions into more developed urban regions.

One potential explanation for the pattern of asymmetric gene flow from rural to urban regions is that diffuse urban areas such as parks, private gardens, and residential neighborhoods represent highly suitable habitats for *Ae. albopictus.* These habitats have a relatively high abundance of resources including oviposition sites, hosts for feeding, and vegetated areas likely resulting in low dispersal out of these areas (Sherpa et al., 2020). The higher genetic differentiation seen in these populations can be attributed to multiple introductions from populations in neighboring rural regions. Some of these urban areas then serve as “hubs” which facilitate gene flow, connecting rural mosquito populations to a multitude of urban habitats that closely fit *Ae. albopictus’* niche. It is possible that the urbanized central regions of Wake County were more highly connected not only to regions within Wake County, but also to other urban centers outside of the county, thus serving as sources of genetic diversity within the observed network. Controlling mosquito dispersal into or out of these sites could potentially help limit gene flow through the overall network of mosquito populations in Wake County.

Our sampling was limited to Wake County and thus did not measure the genetic contribution of *A. albopictus* populations outside of the county boundaries. As noted by Meirmans (2015), biological boundaries do not follow geopolitical jurisdictions, and migration rates likely underestimate the number of immigrants from surrounding areas in the regions along the border of Wake County. Despite the limited sampling, this study was designed to be context-dependent within a geopolitical boundary and provides evidence for complex landscape-driven dispersal. We recommend more comprehensive sampling of urban and suburban areas identified *a priori* in order to further clarify patterns of asymmetrical migration in *Ae. albopictus*.

Certain node statistics, such as degree centrality and closeness centrality, are not the most reliable measure of network connectivity when the number of migrants within a network is high. Koen et al. (2016) found that node degree generally decreased as the number of migrants increased. In a network with few migrants, edge weights were mostly the same and thus were not pruned, resulting in nodes with many edges. As connectivity increased, low-weight edges were pruned, leading to decreasing degree centrality with increasing connectivity, a counterintuitive outcome. As suggested by Koen et al. (2016), we also evaluated a weighted version of degree (mean inverse edge weight), which is a more reliable measure of connectivity that allows us to incorporate the information provided by edge weight.

This study is among a growing body of literature that utilizes network analysis to study gene flow within a network of genetically connected populations (Cross et al., 2018; Dyer and Nason 2004; Garroway et al., 2008; Greenbaum et al., 2016; Jones and Manseau 2022, Koen et al., 2016; Miles et al., 2018; Rodger et al., 2018; Wogan et al., 2024). Population graphs generated based on Dyer & Nason’s conditional independence approach (2004) are able to infer landscape resistance to gene flow at fine spatial scales, particularly when dispersal distances are unknown or when long-distance dispersal events are a possibility (Savary et al., 2021a). This approach estimates the covariance between populations, conditional upon each population’s covariance with the rest of the sampled populations, uncovering a more accurate snapshot of realized dispersal events (Dyer, 2015; Savary et al., 2021a). However, populations may exchange genes over several generations, even if they are not connected by direct dispersal paths (i.e. stepping-stone model). This is especially true for insects that may have multiple generations within a year. Because genetic structure is often shaped by multi-generational dispersal, analyzing population graphs in conjunction with traditional landscape genetic measures such as pairwise *F_ST_* can disentangle the patterns arising from contemporary versus long-term gene flow.

Network analysis can be a powerful tool for ecological studies because it inherently accounts for non-independence and ecological data is often correlated (Fletcher et al., 2011; Jones & Manseau, 2022). While distance-based techniques such as Mantel tests utilize these correlations to test hypotheses of gene flow, these tests tend to have relatively low statistical power and require assumptions of mutation-drift equilibrium (Anderson et al., 2010; Hall & Beissinger, 2014; Legendre & Fortin, 2010; Koen et al., 2016). Landscape genetic analyses often observe poor model fit because genetic distances do not form strongly linear relationships to landscape distances, even with transformations (Diniz-Filho et al., 2013; Shirk et al., 2018). In addition, an inherent weakness of distance-based techniques is that they cannot account for node-based variables that might influence local population dynamics and gene flow (Greenbaum & Fefferman, 2017; Jones & Manseau, 2022; Richardson et al., 2016). This is especially relevant for studying organism dispersal at fine spatial scales, as dispersal behavior can be influenced by habitat quality (node variable) as well as landscape resistance (edge variable). The strength of network analysis lies in its ability to incorporate both node- and edge-based variables, which is not possible with traditional landscape genetics methods such as the Mantel test or multiple regression on distance matrices. Network analysis is a rapidly evolving field—exponential random graph models (ERGMs), which previously only allowed hypothesis testing of binary networks (where a link between nodes is either present or absent), have now been developed for valued networks (Desmarais & Cranmer, 2012; Krivitsky, 2012). While ERGMs have been utilized in ecological contexts to study landscape connectivity (Sánchez-Cano et al., 2024), currently no studies have incorporated genetic data into this approach. Incorporating graph theoretic approaches into landscape genetics can not only help us quantify observed genetic connectivity, but these methods can also be applied to build predictive models, for example evaluating how land use change will influence population connectivity. There are important implications for vector control in public health settings, as a better understanding of how genes move around the landscape will inform the deployment of genetically modified mosquitoes and the pattern and speed of the evolution of resistance to insecticides.

*Aedes albopictus* is a ubiquitous nuisance species and vector of zoonotic disease, and it is important to adopt adaptive, science-based strategies to control its spread. Landscape genetic approaches can be a useful tool for understanding when and for how long control measures will work. Based on network analysis and bidirectional migration estimates, we recommend that mosquito control focus surveillance and management efforts on areas actively growing and urbanizing. Areas identified as likely hubs of movement may be important target sites for an integrated control approach to forestall the spread of resistance or other population characteristics we want to avoid (e.g. ability to transmit a pathogen), or where to release genetically modified mosquitoes for maximal impact. While more research is needed to generalize these recommendations to other locations, fine-scale, place-based studies such as this provide support and context to larger-scale research and are directly applicable to local control agents, connecting science to practice.

## ACKNOWLEDGEMENTS

We would like to thank Anastasia Figurskey, Allison Cousins and Chris Intehar for their assistance in field collection and identification, Emma Wallace for her help with DNA extractions and genomic library preparation, and Paul Labadie for consultation on molecular methods and bioinformatic processing. We also thank the Wake County Department of Health and Human Services for their permission to sample on public lands. This study was supported by the USGS Southeast Climate Adaptation Science Center graduate fellowship awarded to Emily M. X. Reed, the Genetics & Genomics Academy at NC State for the fellowship that supported J.Y. Ding. We thank the Global One Health Academy at NC State for an additional fellowship that supports J.Y. Ding. We would also like to thank the Wynne Innovation Grant from the CAL Dean’s Enrichment Grant Program at NCSU awarded to Martha O. Burford Reiskind that supported the research. This manuscript was greatly improved by three anonymous reviewers.

## DATA ACCESSIBILITY STATEMENT

The genomic data sets and metadata that support this manuscript will be available in MO Burford Reiskind’s Dryad account specific to this publication https://doi.org/10.5061/dryad.2bvq83c2x.

## AUTHORSHIP CONTRIBUTIONS

Jessica Y. Ding: Conceptualization of new analyses, Methodology, Data Collection, Formal analysis, and Writing-original draft. Emily M. X. Reed Conceptualization, Methodology, Data Collection, Formal analysis, and Writing-original draft. Michael H. Reiskind: Conceptualization, Methodology, Writing-reviewing and editing. Martha O. Burford Reiskind: Conceptualization, Methodology, Formal analysis, Writing-reviewing and editing.

---

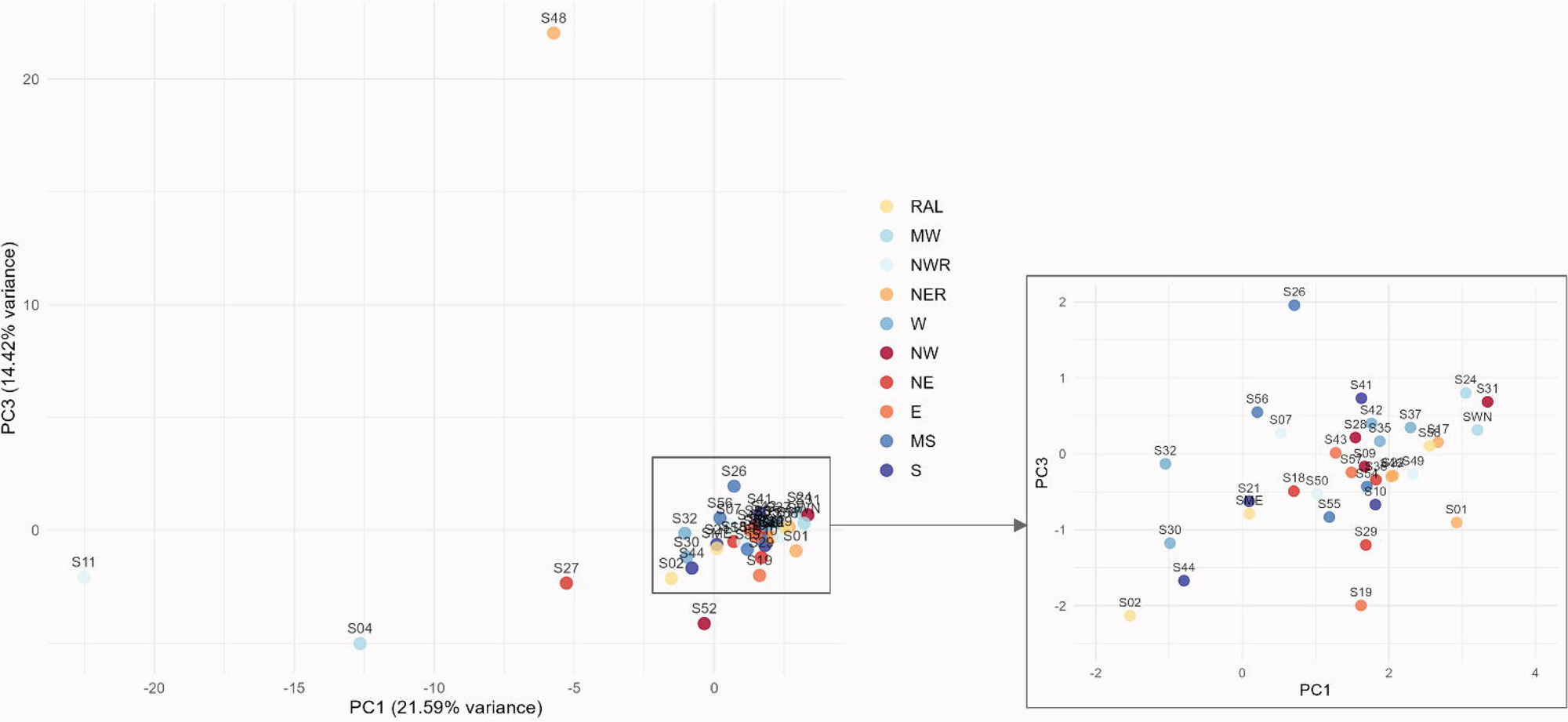

