## Supplemental Tables and Figures for "Fine-scale landscape genomics show asymmetric patterns of gene flow for the invasive mosquito *Aedes albopictus*"

**Supplement**

**Tables:**

**Table S1.** Wake county regions were classified as urban or rural based on distance to major road, % impervious surface, and % forest cover. Regional values were obtained by averaging 100m buffers around each sample site. RAL, NER, NWR, MW, and W had low % forest cover and high % impervious surface and were classified as urban. NW, NE, E, MS, and S had high % forest cover and low % impervious surface and were classified as rural. While S had relatively high % impervious surface compared to other rural regions, we defined it as rural due to its high % forest cover and proximity to other rural regions.

| | | Dist. to major road | % Impervious surface | % Forest cover |
|---|---|---|---|---|
| URBAN | NWR | 89 | 45.6 | 0 |
| | MW | 72 | 42.3 | 0 |
| | RAL | 144 | 36.1 | 1.8 |
| | NER | 287 | 26.3 | 12.2 |
| | W | 284 | 16.2 | 30.4 |
| RURAL | MS | 602 | 5.3 | 32.5 |
| | E | 502 | 6.7 | 34.2 |
| | S | 454 | 17.9 | 41.4 |
| | NE | 411 | 6.7 | 47.3 |
| | NW | 647 | 1.8 | 81.0 |

**Table S2.** Number of individuals per site. Includes total number collected, number of individuals sequenced, and number of individuals retained for analysis for each site. Rows highlighted in grey indicate sites that were excluded from analysis because there were too few individuals.

| Site ID | Total # collected | # extracted | # sequenced | genind.unfiltered | genind.FILTERED |
|---|---|---|---|---|---|
| **S01** | 62 | 7 | 6 | 6 | 6 |
| **S02** | 7 | 7 | 6 | 6 | 6 |
| **S03** | 4 | 4 | | | |
| **S04** | 16 | 10 | 6 | 5 | 5 |
| **S06** | 3 | 3 | | | |
| **S07** | 75 | 11 | 6 | 6 | 6 |
| **S08** | 0 | | | | |
| **S09** | 21 | 12 | 16 | 8 | 8 |
| **S10** | 6 | 6 | 6 | 6 | 6 |
| **S11** | 25 | 10 | 6 | 5 | 4 |
| **S12** | 2 | 2 | | | |
| **S14** | 3 | 3 | | | |
| **S15** | 5 | 5 | | | |

| | | | | | |
|---|---|---|---|---|---|
| **S17** | 132 | 11 | 16 | 8 | 7 |
| **S18** | 14 | 8 | 6 | 6 | 6 |
| **S19** | 8 | 7 | 6 | 6 | 5 |
| **S20** | 3 | 3 | | | |
| **S21** | 26 | 15 | 10 | 9 | 9 |
| **S22** | 323 | 7 | 6 | 6 | 6 |
| **S23** | 6 | 6 | | | |
| **S24** | 24 | 13 | 6 | 6 | 6 |
| **S25** | 19 | 9 | 6 | 5 | |
| **S26** | 7 | 7 | 6 | 6 | 6 |
| **S27** | 6 | 6 | 6 | 6 | 4 |
| **S28** | 18 | 9 | 6 | 6 | 6 |
| **S29** | 51 | 11 | 6 | 6 | 6 |
| **S30** | 12 | 9 | 7 | 7 | 7 |
| **S31** | 14 | 13 | 16 | 7 | 6 |
| **S32** | 36 | 13 | 7 | 7 | 7 |
| **S33** | 12 | 11 | 1 | | |
| **S34** | 5 | 5 | | | |
| **S35** | 16 | 8 | 6 | 6 | 6 |
| **S36** | 8 | 8 | 8 | | |
| **S37** | 28 | 8 | 6 | 6 | 6 |
| **S38** | 13 | 11 | 8 | 8 | 7 |
| **S39** | 5 | 5 | | | |
| **S40** | 8 | 7 | 6 | | |
| **S41** | 176 | 8 | 6 | 6 | 6 |
| **S42** | 19 | 15 | 16 | 8 | 8 |
| **S43** | 30 | 10 | 7 | 7 | 7 |
| **S44** | 44 | 7 | 6 | 6 | 5 |
| **S45** | 0 | | | | |
| **S46** | 49 | 8 | 6 | 6 | 6 |
| **S47** | 11 | 8 | 7 | 6 | |
| **S48** | 41 | 7 | 6 | 5 | 5 |
| **S49** | 59 | 7 | 6 | 6 | 6 |
| **S50** | 58 | 7 | 7 | 7 | 7 |
| **S51** | 5 | 5 | | | |
| **S52** | 14 | 7 | 6 | 6 | 5 |
| **S53** | 1 | | | | |
| **S54** | 25 | 8 | 8 | 8 | 8 |
| **S55** | 41 | 7 | 7 | 7 | 7 |
| **S56** | 37 | 7 | 7 | 7 | 6 |
| **S57** | 10 | 10 | 8 | 8 | 8 |

| | | | | | |
|---|---|---|---|---|---|
| **S58** | 175 | 12 | 16 | 8 | 8 |
| **S60** | 5 | 5 | | | |
| **SME** | 110 | 9 | 7 | 7 | 7 |
| **SOO** | 5 | 5 | | | |
| **SPV** | 11 | | | | |
| **SVD** | 52 | 10 | 7 | 7 | |
| **SWN** | 85 | 8 | 6 | 6 | 5 |
| **TOTAL** | **2086** | **460** | **336** | **274** | **245** |

**Table S3.** Summary of bioinformatic processing and single nucleotide polymorphism (SNP) filtering. Minimum minor allele frequencies (MAF) is set at 0.01.

| | Denovo Pipeline (STACKS) | Populations Pipeline (STACKS) | MAF (Plink) | GENO (Plink) | MIND (Plink) | HWE (Plink) | Final Dataset (RStudio) |
|---|---|---|---|---|---|---|---|
| **Individuals** | 336 | 289 | 289 | 289 | 274 | 274 | 245 |
| **Variants** | 3096027 | 183753 | 169713 | 16066 | 16066 | 11305 | 4,013 |

**Figures:**

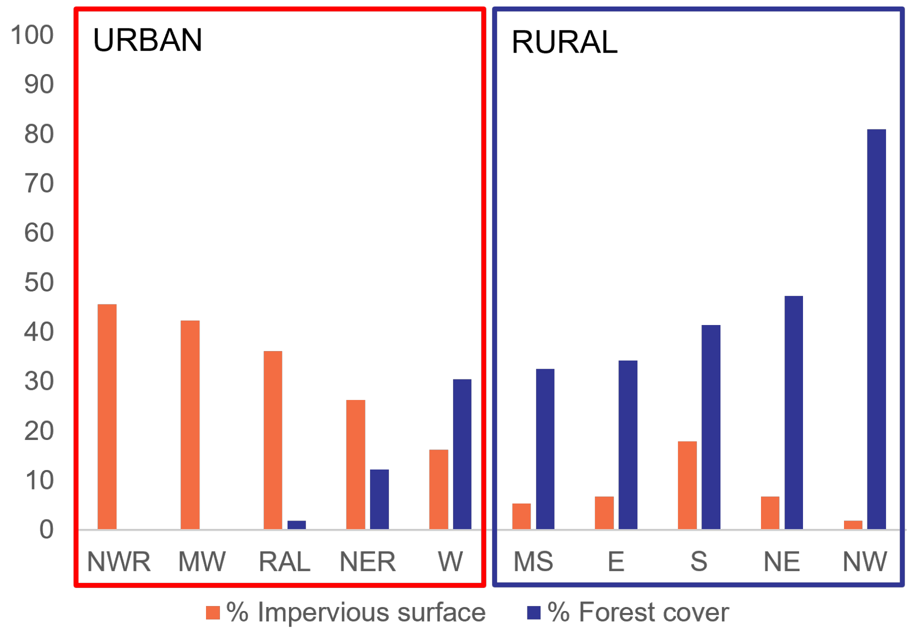


**Figure S1.** Average % impervious surface and % forest cover by Wake County region. Regional means were obtained by averaging 100m buffers around each sample site. NWR, MW, RAL, NER, and W regions were classified as urban, while MS, E, S, NE, and NW were classified as rural.

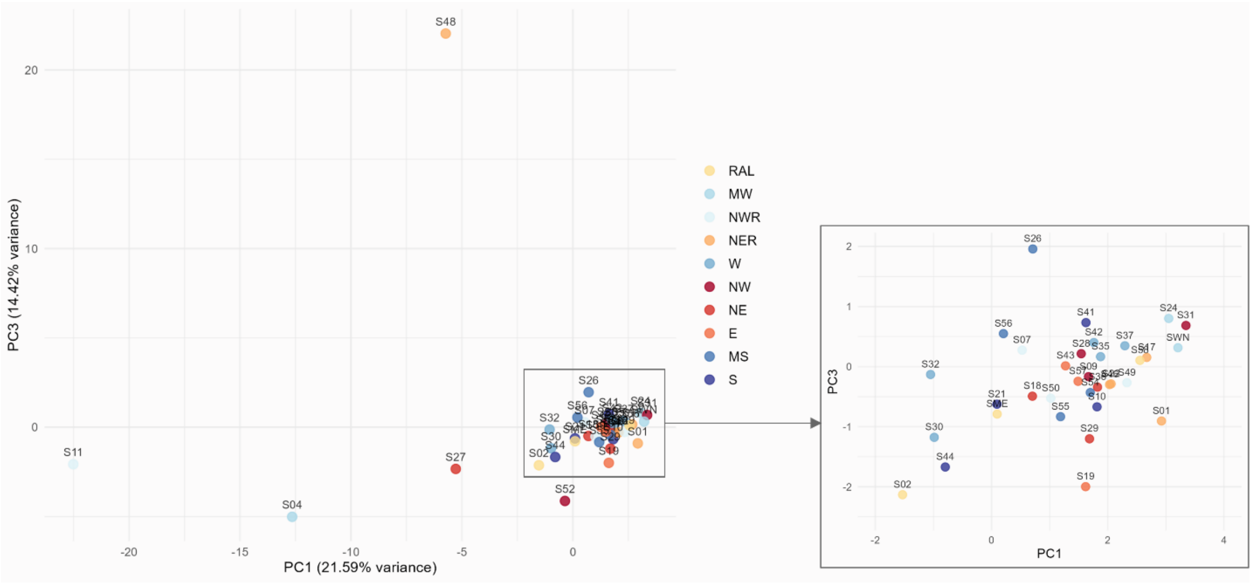


**Figure S2.** Population-level PCA with PC1 and PC3 on the *x* and *y* axes, respectively. Points are colored according to Wake County regions (see **Figure 2**). PC1 accounts for 21.59% of the variance, while PC3 accounts for 14.42% of the variance. The majority of sites again form a single cluster with no clear clustering patterns by region. S04, S11, S27, S48, and S52 demonstrate greater differentiation from the main cluster, with S52 located in a rural region and the other sites located in urban regions.
